# Multi-omics analysis reveals critical cis-regulatory roles of transposable elements in livestock genomes

**DOI:** 10.1101/2023.08.17.553652

**Authors:** Chao Wang, Bowen Lei, Yongzhou Bao, Zhen Wang, Choulin Chen, Yuanyuan Zhang, Shenghua Qin, Tao Sun, Zhonglin Tang, Yuwen Liu

## Abstract

As a major source of genetic and regulatory variation in their host genome, transposable elements (TEs) have gained a growing interest in research on humans and model organisms. In this species, integrative analysis of multi-omics data has shed light on the regulatory functions of TEs. However, there remains a notable gap in our understanding of TEs in domesticated animals. we annotated TEs in the genomes of pigs, cattle, and chickens, respectively, and systematically compared the genome distributions and amplification patterns of TEs across these three species. Furthermore, by integrating multi-tissue RNA-seq, ATAC-seq, and histone modification ChIP-seq data, we explored the expression atlas of TEs and their contribution to cis-regulatory elements (CREs) in different tissues of the three species. Most importantly, we developed a novel computational framework that revealed TE-mediated gene regulatory networks (TE-GRNs) underlying tissue-related biological processes. To demonstrate the power of this approach, we applied our framework to analyze liver tissues across the three different species. Overall, our research provides novel insights into the regulatory functions of TEs in livestock animals and highlights a computational framework to uncover TE-GRNs in various biological contexts.

## Background

Making up between 4% to 60% of the vertebrate genomes [1], TEs are a type of repetitive DNA sequences that can replicate and change their positions within a host genome. Due to differences in transposition mechanisms, TEs can be classified into two classes: DNA transposons and retrotransposons. DNA transposons primarily exploit a “cut and paste” mechanism to move from one genomic location to another, while retrotransposons use a “copy and paste” mechanism, by being first transcribed into RNA and then reverse transcribed to DNA, and then inserted to a new location in the genome [2–4]. Based on sequence composition, retrotransposons could be further divided into SINE, LINE, and LTR classes. TEs were initially considered to be “junk DNA” which only serve to invade the genome through their transposition ability. However, due to this replicability and mobility, TEs are also an extensive source of mutations and genetic polymorphisms in genomic regions that gradually manifest as functional [5–7]. Thus, by creating or deleting these important functional DNA sequences, TEs exhitibed crucial roles in driving genome evolution and gene regulation [8–10]. For example, TEs can directly influence the coding sequences of genes [11–14], and provide raw material for the emergence of non-coding RNAs, such as LncRNAs [15], microRNAs [16] and other small RNAs [17]. They can also function as the building blocks of DNA CREs, such as promoters and enhancers [18–20], insulators [21], and silencers [22].

The diverse mechanisms by which TEs can exert influence on the genome have sparked significant interest in the development of versatile tools to study them. These include TE annotation tools, such as ReaptMasker [23], REPET [24], phRAIDER [25], RepeatExplorer [26], dnaPipeTE [27], and DeepTE [28], as well as TE expression quantification tools, such as Telescope [29], TEtranscripts [30], and SQuIRE [31]. Beyond annotating TEs and quantifying their expression, the functional interpretation of TEs were greatly strengthened by functional genomics projects such as ModENCODE for model organisms [32], ENCODE for humans [33]. These studies generated multi-omics functional genomics data that predict the cis-regulatory activity of DNA sequences, thus shedding light on how TEs rewire the transcriptional regulatory networks by influencing CREs. For instance, Pehrsson et al. integrated data from the Roadmap Epigenomics Project to analyze the contribution of different TE classes to CREs predicted by multiple epigenomic marks across human anatomy and development [34]. Choudhary et al. used multi-species 3D genome data to identify TE families and subfamilies that impact lineage-specific chromatin structures during the evolution of gene regulation [35]. Chang et al. performed a systematic analysis of TE age and genomic distribution in zebrafish, and used both bulk and single-cell transcriptomic data to explore TE expression patterns during development [36]. Lee et al. integrated multi-omics data to unravel that TEs significantly contribute to diverse tissue-specific CREs and in zebrafish. They also showed that TEs can drive the formation of promoters and interfere with gene transcription [37]. By integrative analysis of multi-omics data with TE annotation, these studies provide possible mechanisms by which TEs might impact complex diseases and traits.

In comparison to humans and model organisms, the study of TEs in livestock has received considerably less attention. As a result, the extent to which TEs influence economic traits in livestock remains largely unexplored. While recent studies in the field of livestock functional genomics have generated mutli-tissue CRE profiles in various domesticated animals [38–42], how TEs contributed to the formation of these CREs and their impact on the rewiring of gene regulatory networks (GRNs) in different tissues is not well understood.

In this study, we annotated TEs in the genomes of pigs, cattle, and chickens, respectively. In these species, we comprehensively analyzed TEs’ genomic distribution, age, expression activity, and epigenetic regulation patterns by combinging ChIP-seq, ATAC-seq, and RNA-seq data from multiple tissues. To elucidate how TEs function as CREs to regulate biological processes, we also developed a computational framework to construct TE-GRNs. Overall, our study established a foundation for future work on analyzing how TEs contribute to the genetic basis of economically imporatant traits in livestock.

## Result

### The genomic landscape of TEs in the genomes of pigs, cattle and chickens

We annotated TEs in three major livestock species, namely chickens (Gallus gallus), pigs (Sus scrofa), and cattle (Bos taurus), using the RepeatMasker software. In total, 3,953,666, 4,984,795, and 322,047 TEs were identified in pigs, cattle, and chickens, respectively. TEs account for approximately 9.6% of the chicken genome and a significantly higher proportion in the pig genome (43.4%) and cattle genome (47.5%) (Fig. 1A). This is consistent with previous research that demonstrated a positive correlation between the TE contents and the genome size [43–45]. We further explored the genome coverage of different TE classes across the three species and found that the LINE class had the highest genome coverage in all three species, followed by the SINE class in the genomes of pigs and cattle. However, the genome coverage of the SINE class in the chicken genome was significantly lower (Fig. 1B). Additionally, we investigated the relative abundance of each TE class among all annotated TEs in the genome. In pigs and cattle, SINE and LINE retrotransposons were found to be the most abundant, whereas in chickens, LINEs were the prevailing retrotransposons, with a considerably lower proportion of SINEs compared to pigs and cattle. Notably, In pigs and cows, although SINE has a higher transposon count than LINE, LINE has a higher genome coverage than SINE (Fig. 1B-C).

**Figure 1.**
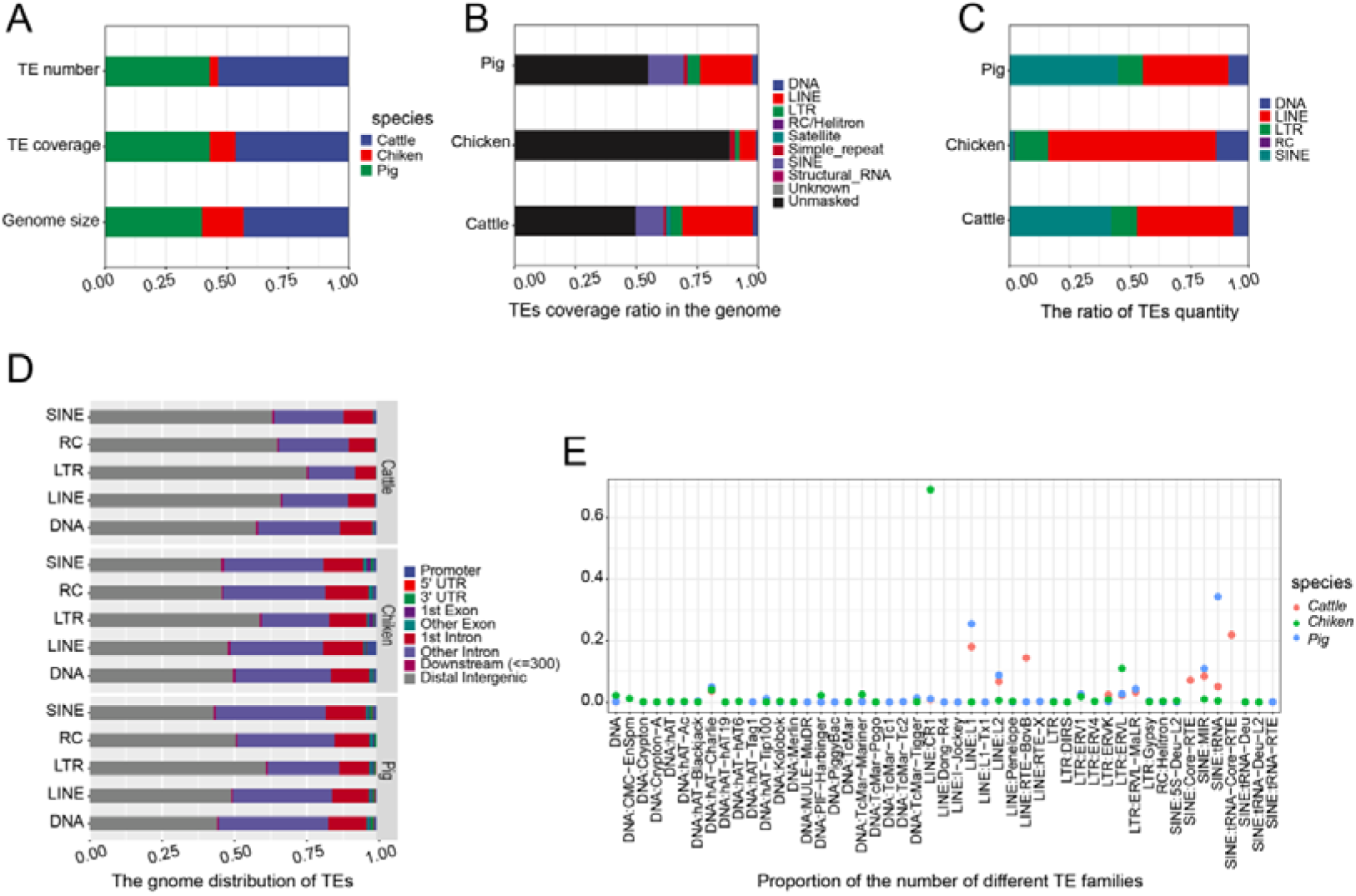
TE annotation and genomic landscape across pigs, cattle and chickens. (**A)** TE quantity, genomic coverage, and genome size proportion distribution in the three species. (**B)** Comparison of genomic coverage of different TE classes in the three species. (**C)** Proportion of different TE classes among all annotated TEs in the three species. (**D)** Genomic distribution of different TE classes in the three species. (**E)** Proportion of different TE families among all annotated TEs in the three species.

Depending on their genomic insertion sites, TEs can potentially act as CREs to regulate gene expression or directly affect protein-coding sequences of genes. We analyzed the genomic distribution of different TE classes and families and found that, in all three species, TEs tended to insert into distal intergenic and intronic regions (Fig. 1D and Additional file 1: Fig. S1A-C), suggesting that the majority of TEs might serve as potential CREs. Additionally, LTR retrotransposons exhibited a higher preference in the non-coding genomic regions compared to other types of TEs (Fig. 1D).

To further understand the contribution of different TE families to genome composition, we analyzed the proportion of each TE family in the total annotated TEs. We found that each species has species-specific TE families that substantially outnumbered other TE families. For instance, in chickens, the LINE/CR1 family exhibits a notably higher proportion compared to other retrotransposon families. In pigs, the SINE/tRNA family displays a particularly elevated proportion. On the other hand, in cattle, the families of SINE/Core-RTE, SINE/tRNA-Core-RTE, and LINE/RTE_BovB demonstrate significantly higher proportions (Fig. 1E). This result indicates that a TE family may have different functional importance and thus were subject to species-specific selection during evolution.

Overall, we successfully annotated TEs in the genomes of three major livestock species, namely chicken, pig, and cattle, and compared the similarities and differences in their genomic distribution and abundance, providing a solid foundation for further research on how TEs function in the livestock genomes.

### Amplification patterns of TEs differ among different livestock species

The abundance and genomic distribution of TEs are closely linked to the evolutionary dynamics of TEs. Therefore, we investigated the transposition rate of TE classes at different time points during species evolution in pigs, cattle, and chickens. Our findings revealed that each species exhibited distinct bursts of TE amplification. Although LINEs and SINEs were more abundant than other TEs in pigs and cattle, their burst patterns differed between the two species. Pigs experienced two bursts, while cattle had three (Fig. 2A-B). In chickens, both SINEs and LINEs had only one round of burst, but LINEs maintained high levels of amplification while SINEs lost their ability to amplify in the modern chicken genome (Fig. 2C). Moreover, DNA TEs had one round of burst in pigs and cattle but two in chickens. On the other hand, LTRs displayed only one round of burst in all three species (Fig 2A-C). We further investigated the transposition rate of TE families within each class in the three specises. Our analyses showed that, within a given TE class, distinct TE families exhibited varying patterns of amplification (Fig. 2D), as evidenced by differences in the timing and amplitude of burst. Accordingly, only some TE families are still active in the modern genomes of pigs, cows, and chickens. In the SINE class, all transposon families in chickens have lost their transposition activity, while Core-RTE and tRNA families are still active in cattle and pigs, respectively. Within the LINE class, RTE-BovB and L1 families in cattle, L1 family in pigs, and CR1 family in chickens are still capable of transposition. In the LTR class, ERVK and ERV1 families of cattle, and ERVK and ERVL families of chickens, still exhibit transposition activity, while all LTR families of pigs have lost their activity almost entirely. Notably, all DNA families have lost transposition activity in the modern genomes of all three species.

**Figure 2.**
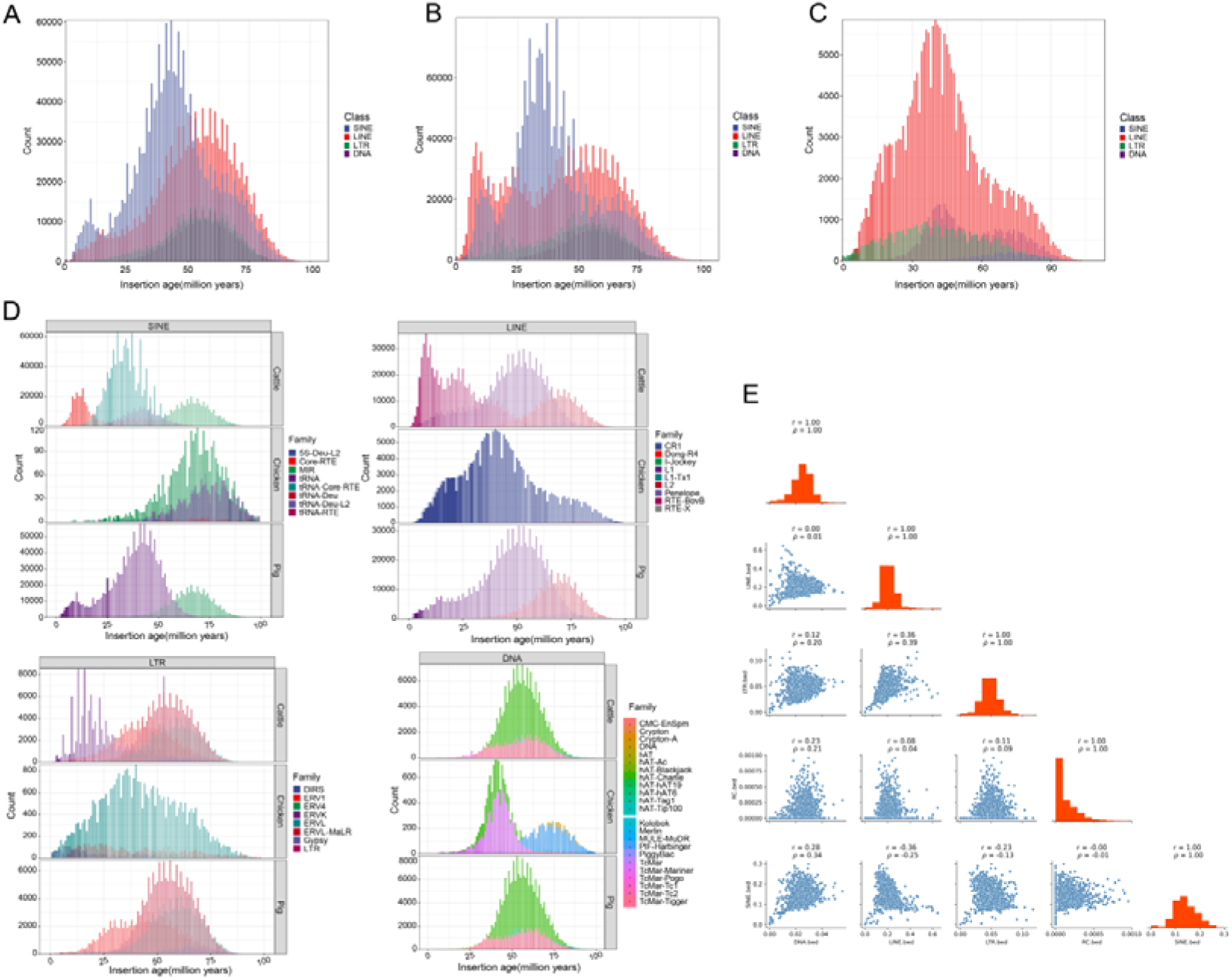
Age distribution and genomic distribution correlation of different types of TEs in pigs, cattle and chickens. (**A-C)** Age distribution of different TE classes in pig(**A**), cattle (**B**), and chicken (**C**). (**D)** Age distribution of different TE families of specific TE classes in three species. (**E)** The genomic distribution relationship of different TE classes by calculating the density of each TE class (as genome sequence coverage) in non-overlapping 2-Mb windows along the genome and calculated the pairwise correlation between different TE classes.

To explore the genomic distribution relationship of different TE classes, we adopted the approach used in a previous study, which calculated the density of each TE class (as genome sequence coverage) in non-overlapping 2-Mb windows along the genome and then analyzed the pairwise correlation between different TE classes[36]. Using this method, we revealed the correlation of genomic distribution between TE pairs of interest. Specifically, we found that LTR and LINE densities are positively correlated in all three species (Fig. 2E, and Additional file 1: Fig. S1D-E). In addition, there are species-specific correlation patterns between TE classes. For example, in pigs and cattle, SINE density is negatively correlated with LINE density (Fig. 2E, and Additional file 1: Fig. S1D). In chickens, LTR density exhibits a high negative correlation with SINE density, whereas LTR density is positively correlated with LINE and LTR densities, respectivelly (Additional file 1: Fig. S1E).

Overall, we systematically analyzed the amplification patterns and correlation of genomic density distribution for different TE classes in the three species, which may shed light on the significant events that these species have experienced throughout their evolutionary history.

### TEs play an important role in regulating tissue-specific chromatin accessibility

Open chromatin regions (OCRs) are exposed in intracellular environment, vulnerable to transcription factor (TF) binding and associated with cis-regulatory activity affecting gene exression. Using ATAC-seq data from five production trait-related tissues (muscle, fat, lung, spleen, and liver) in the three livestock species, we explored the chromatin accessibility of different TE classes. The SINE class was significantly enriched for open chromatin signals in every tissue of the three species, while the LINE and DNA classes were significantly depleted. An intriguing observation is the depletion of chromatin-accessible signals in LTR across all tissues in chickens. In contrast, pigs and cattle exhibit relatively high levels of enrichment of chromatin-accessible signals attributed to LTR (Fig. 3A). These results suggest that SINEs have the potential to act as CREs in all three species, while the cis-regulatory potential of LTRs are specific in pigs and cattle.

**Figure 3.**
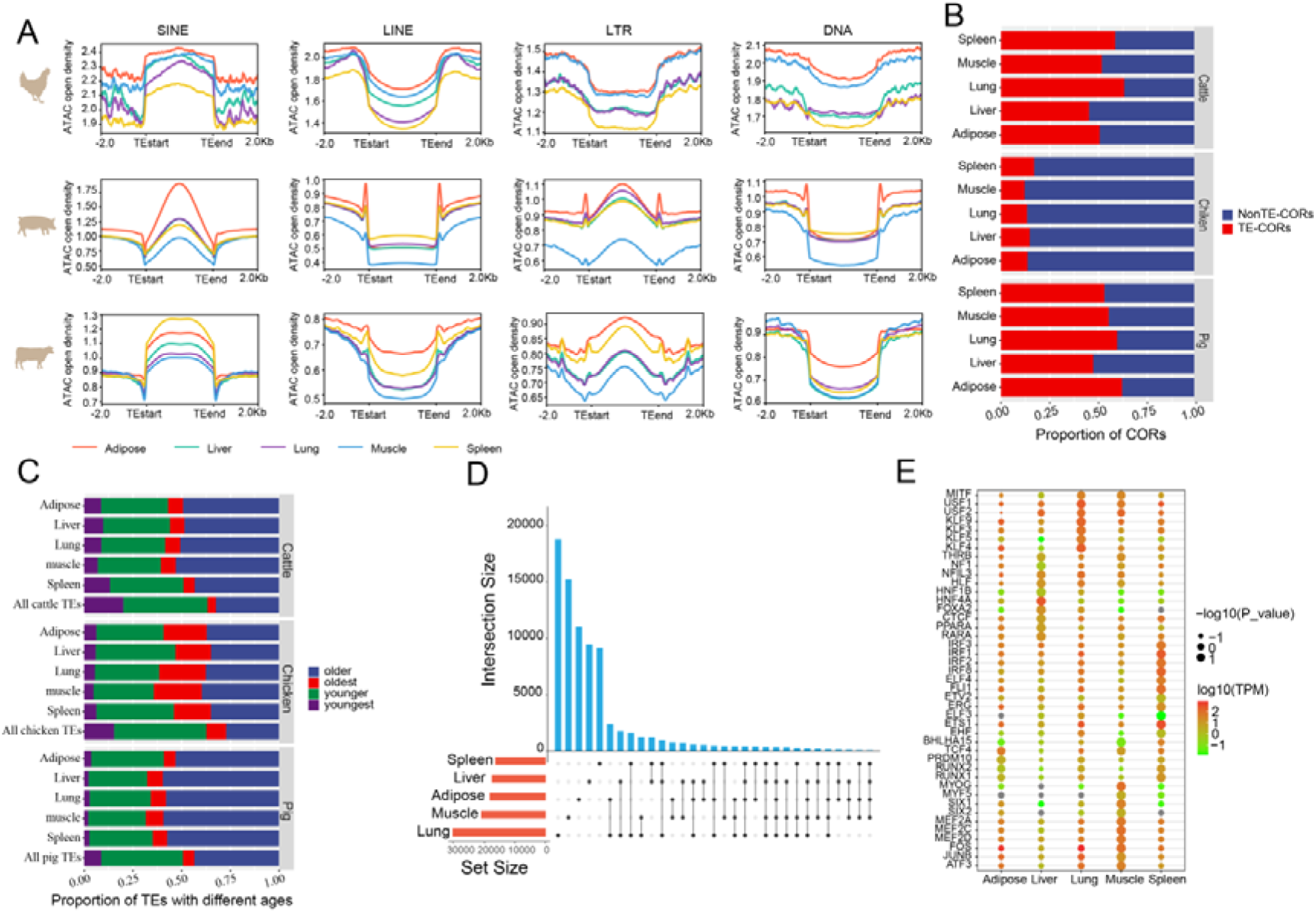
The effect of TEs on OCRs in various tissues of pigs, cattle, and chickens. (**A)** The reads density distributions of chromatin accessibility of four major TE classes across different tissues in three species. (**B)** The distribution of the proportion of TE-driven and non-TE-driven OCRs across different tissues in three species. (**C**) The age distribution of OCRs-residing TEs in different tissues across three species. (**D)** The bar plot shows the number of chromatin-accessible tissue-specific or shared TEs in pig across different tissues. The red bar represents the total number of TEs with chromatin accessibility features in that tissue, and the blue bar represents the number of TEs with chromatin accessibility features shared among specific tissues. (**E)** Bubble plot showing significant enrichment of TF motifs in chromatin-specific accessible TEs in each tissue of pig, along with the expression levels of the corresponding TFs in that tissue. Select the top 20 significantly enriched TFs (Q-value < 0.01) in each tissue and display only the TFs that are expressed in that tissue.

To investigate the impact of TEs on OCRs in different tissues, we analyzed the number of OCRs overlapping TE insertions in each tissue. We defined OCRs with TE insertions as TE-driven OCRs (TE-OCRs). In pigs and cattle, TE-OCRs accounted for around 50% of all OCRs, while in chickens, they constituted only approximately 15% of OCRs (Fig. 3B). These findings suggest that the degree to which TEs contributed to creating CREs substantially varies between chickens and the other two species. To further explore the impact of different TE classes on OCRs in different tissues, we analyzed the proportion of the four major TE classes within TE-OCRs. Across tissues, we found consistent proportions of all four TE classes in TE-OCRs within each species. When compared to all annotated TEs, our analysis revealed a significant depletion of LINEs and an overrepresentation of LTRs within TE-OCRs across all tissues in pigs and cattle. In chickens, LTRs showed a depletion within TE-OCRs, while SINEs exhibited an elevated presence across all tissues (Additional file 1: Fig. S4A). These findings suggest that the contribution of different TE classes to functional CREs is conserved across tissues but varied across species.

Furthermore, we classified TE into four distinct types based on their estimated age: youngest (0-25MYa), younger (25-50MYa), older (50-75MYa), and oldest (75-100MYa). This categorization allows for exploring the enrichment patterns of TEs of varying ages within TE-OCRs across diverse tissues. Using all annotated TEs as the background, we found a significant enrichment of old TEs in TE-OCRs across different tissues in the three species, while young TEs were depleted (Fig. 3C). This finding indicates that the preference for older TEs in forming CREs is not limited to a specific species but is, in fact, a cross-species phenomenon. Thus, our analysis underscores the evolutionary significance of older TEs in shaping the regulatory landscape across diverse organisms.

Moreover, we were interested in understanding the extent to which TEs contribute to tissue-specific CREs, as these CREs are typically involved in tissue-specific biological processes[39]. Across all the three species, we observed a clear pattern that the majority of TEs within TE-OCRs exhibit tissue specific chromatin accessibility (Fig. 3D, Additional file 1: Fig. S2A-B).We hypothesized that these tissue-specific TEs contributed to the formation of tissue-specific CREs, which recruit binding of TFs that play crucial roles in maintaining tissue-specific biological processes. To test this hypothesis, we performed motif enrichment analysis for tissue-specific TEs. After selecting the top 20 enriched TF motifs in tissue-specific TEs, we only retained TFs that were significantly enriched (Q-value < 0.01) and expressed in the corresponding tissue. Surprisingly, we found that tissue-specific TEs in all three species significantly enriched for DNA binding motifs of TFs that are closely relevant to tissue development and highly expressed in the corresponding tissues (Fig. 3E, and Additional file 1: Fig. S2C-D). This result highlights the important role in regulating tissue-related biological processes of tissue-specific TEs with tissue-specific chromatin accessibility.

### TEs affect the epigenetic states of CREs

In order to comprehensively investigate the impact of TEs on specific types of CREs, we classified CREs into promoters, enhancers, and candidate silencer elements (CSEs) based on their epigenetic states (Fig. 4A, detailed in the “Methods” section). Based on the identification methods of these CREs, we successfully constructed atlas of three types of CREs across five tissues in the three species (Fig. 4B, left panel; Additional file 2: Table S1). To ensure that the identified CSEs can truly be classified as a specific type of element with a unique pattern of nucleotide compositions, we used deep learning to test whether CSEs could be distinguished from other sequences solely by their DNA sequences. The model we built achieved good predictive ability for CSEs, as demonstrated by high AUC values in chickens (0.759), pigs (0.850), and cattle (0.857), respectively (Additional file 1: Fig. S3A-C). As expected, the model could also well predict enhancers and promoters. The performance of the model in predicting chicken CREs exhibits a slight decrease, potentially attributed to the limited amount of training data available for chickens.

**Figure 4.**
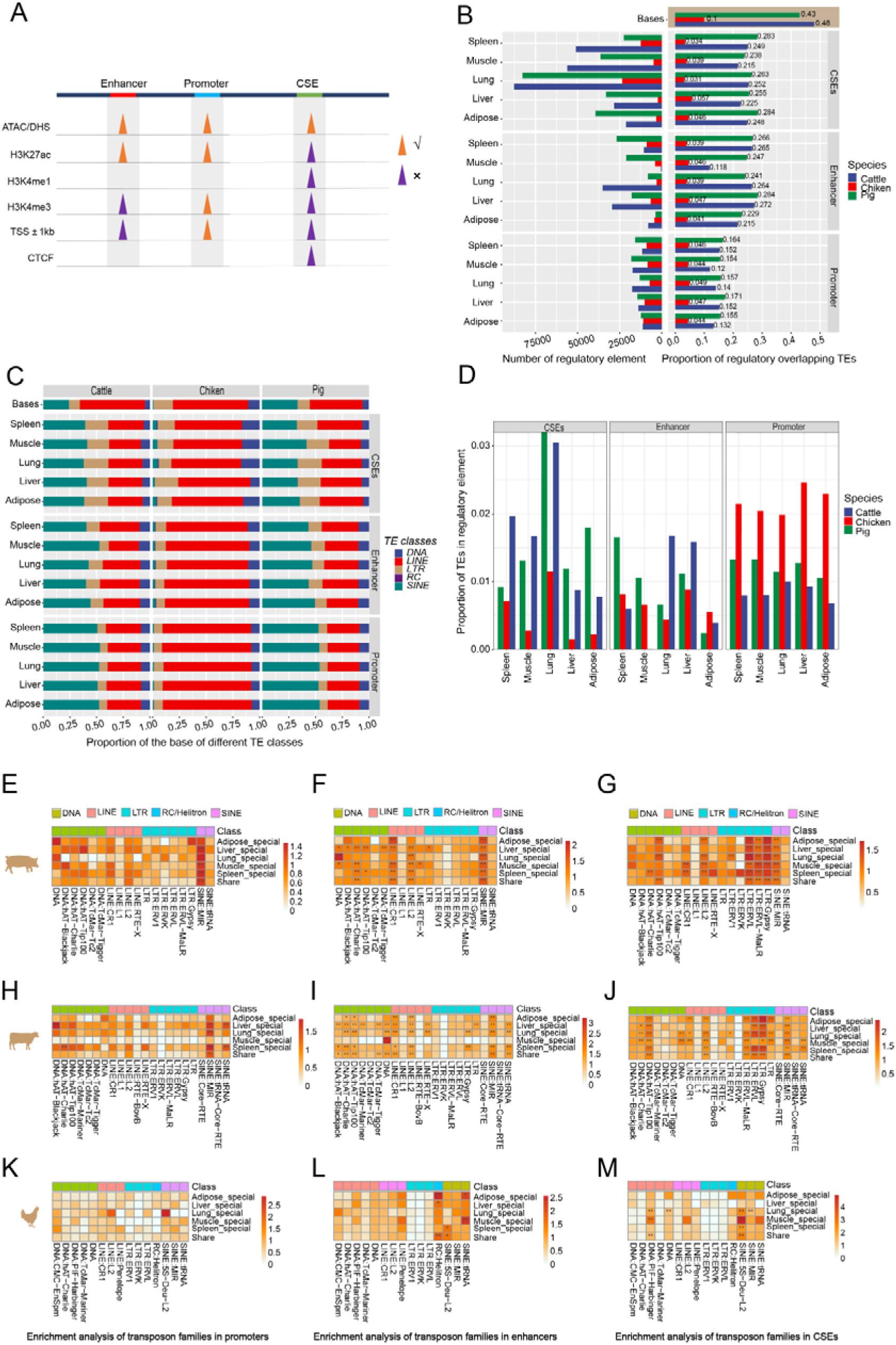
The contribution of TEs to different CREs across different tissues and three species. (**A)** Diagram of CREs categorization. √ indicates the presence of the epigenetic mark, × indicates the absence of the epigenetic mark. (**B)** Number of three types of regulatory elements identified in different tissues of three species (left panel). The base proportion of CREs within TEs across five tissues in three species (right panel). The beige block shows the coverage of TE bases in the genome of three species. (**C)** The distribution of base proportion of different TE classes within specific regulatory elements across different tissues of three species. Base proportion distribution of different TE classes in all annotated TEs of the genome as background. (**D)** The proportion of TEs with specific regulatory element states among all annotated TEs in the genome across different tissues of three species. (**E-M)** TE family enrichment in different types of CREs across the three species. (*) FDR < 0.05 and (**) FDR < 0.01 after correcting for multiple testing using the Bonferroni method.

To investigate the extent to which CREs were constructed on TEs, we analyzed the proportion of TE bases within CREs across five tissues in the three species (Fig. 4B, right panel). With the TE base coverage in the genome as the background, we observed a depletion of TEs in CREs across different tissues in the three species. This pattern could be explained by the defense mechanisms of host genome against transposon insertion, which prevent the insertion of TEs into important functional DNA regions, such as CREs, as a way to minimize deleterious impact on the host’s fitness[46]. Across tissues within a species, the TE base proportion of each CRE type present is similar. However, there were variations in the TE base proportions among different CRE types within a species. In pigs and cattle, enhancers and silencers showed a higher proportion of TEs compared to promoters. In chickens, the TE base proportion of CREs was notably smaller, aligning with the lower overall genome coverage of TEs in the chicken genome (Fig. 4B, right panel). To further investigate how CREs were constructed on different TE classes, we investigated the contributions of different classes of TEs to CREs (Fig. 4C). Using the base proportions of different TE classes in the genome as the background, we found notable variations in the preference of different TE classess within CREs in pigs, chickens and cattle. Specifically, in pigs and cattle, LINEs exhibited depletion across all types of CREs, whereas SINEs displayed enrichment within CREs, except for pig CSEs. Additionally, both pig and cattle CSEs exhibited enrichment of LTRs. In chickens, DNA elements were enriched in CSEs, while LINEs were enriched in enhancers and promoters (Fig. 4C).

We also examined the percentage of TEs that overlapped with promoters, enhancers and silencers, respectively (Fig. 4D). Our analysis revealed that the proportion of TEs overlapping enhancers and silencers varied considerably across tissues in all three species, while the proportion of TEs overlapping promoters remained relatively stable. Notably, in chickens, the proportion of TEs overlapping enhancers and silencers was relatively lower compared to that of pigs and cattle. However, the proportion of TEs overlapping promoters in chickens is higher. These findings suggest that TEs in chickens may have a greater propensity to directly engage in mediating promoter function. Moreover, we observed that the proportion of different TE classes overlapping each CRE type were stable across tissues within each species. However, notable differences were observed in the CRE-overlapping proportion of some TE classes across species (Additional file 1: Fig. S4B).

To further explore the impact of different TE families on different types of CREs, we categorized the TEs by family, and the CREs by tissue-specificity. Subsequently, we utilized permutation tests to evaluate the enrichment of different TE families in CREs specifically active in various tissues. In all three species, we found that fewer TE families were enriched in tissue-specific promoters than in enhancers and CSEs (Fig. 4E-M). Particularly, the chicken promoters did not enrich for any type of TE families (Fig. 4K). These results suggest that TEs made less contribution to promoters than to other CREs. Additionally, in pigs and cattle, we found that LTR families were more enriched in CSEs than in enhancers of different tissues (Fig. 4F,G,I,J), indicating that LTRs generally have a greater impact on CSEs than enhancers. An important observation is that we found a significant enrichment of LTR families in the CSEs of various tissues in pigs and cattle, but not in chickens (Fig. 4G,J,M). This finding provides a plausible explanation for the distinct chromatin accessibility patterns observed in LTRs among pig, cattle, and chicken tissues. Notably, LTRs exhibit a considerably higher level of accessibility in pig and cattle tissues, while demonstrating comparatively lower accessibility in chicken tissues (Fig. 3A). To further investigate why LTRs have a greater impact on CSEs activity in pig and cattle tissues compared to chickens, we performed a motif enrichment analysis of LTR DNA sequences within the CSEs of the three species by HOMER software. After selecting the top 20 significantly enriched TFs (Q<0.01) that also have orthologues in that species, we identified four TFs (MITF, USF1, CLOCK, MNT) that are shared between pigs and cattle, while no TFs are shared between chickens and either pigs or cattle (Additional file 1: Fig. S4C). Importantly, among the four TFs enriched in LTRs of CSEs in pigs and cattle, USF1 has been reported to display transcriptional repressor activity [47, 48]. This findings proposed possible mechanism that LTRs might affect the activity of CSEs specifically in cattle and pigs by recruiting repressive TFs.

Altogether, Our analyses offer a theoretical framework for assessing the significance of transposable elements (TEs) in species-specific and tissue-specific cis-regulatory elements (CREs). This framework facilitates further investigation into how TEs reconfigure transcriptional regulatory networks in diverse tissues and contribute to the development of complex traits.

### Tissue-specific expression of TEs are involved in tissue-related biological processes

TEs not only function as CREs, but they can also regulate gene expression in trans through their transcription. In order to investigate the potential impact of TE expression on biological processes, we downloaded transcriptome data from five tissues of chickens, pigs and cattle, each with two biological replicates. We then utilized SQuIRE software to construct TE expression atlas across tissues within each species [31]. Our study defined a TE as transcriptionally detectable in a tissue when its average FPKM in that tissue is greater than 1. Interestingly, we found that although there were significant differences in the number of detectable TEs among each species and across different tissues, the cross-tissue fluctuation pattern in the number of detectable TEs was similar among pigs, cattle, and chickens, with the spleen having the highest detectable TEs and muscle the lowest (Fig. 5A). Subsequently, we analyzed the proportion of detectable TE classes in different tissues (Additional file 3: Table S2A-C). Compared to all annotated TEs, SINEs were preferentially expressed in each tissue of the three species. However, LINEs still comprised the largest proportion of detectable chicken TEs due to their predominant presence in the chicken genome (Fig. 5B).

**Figure 5.**
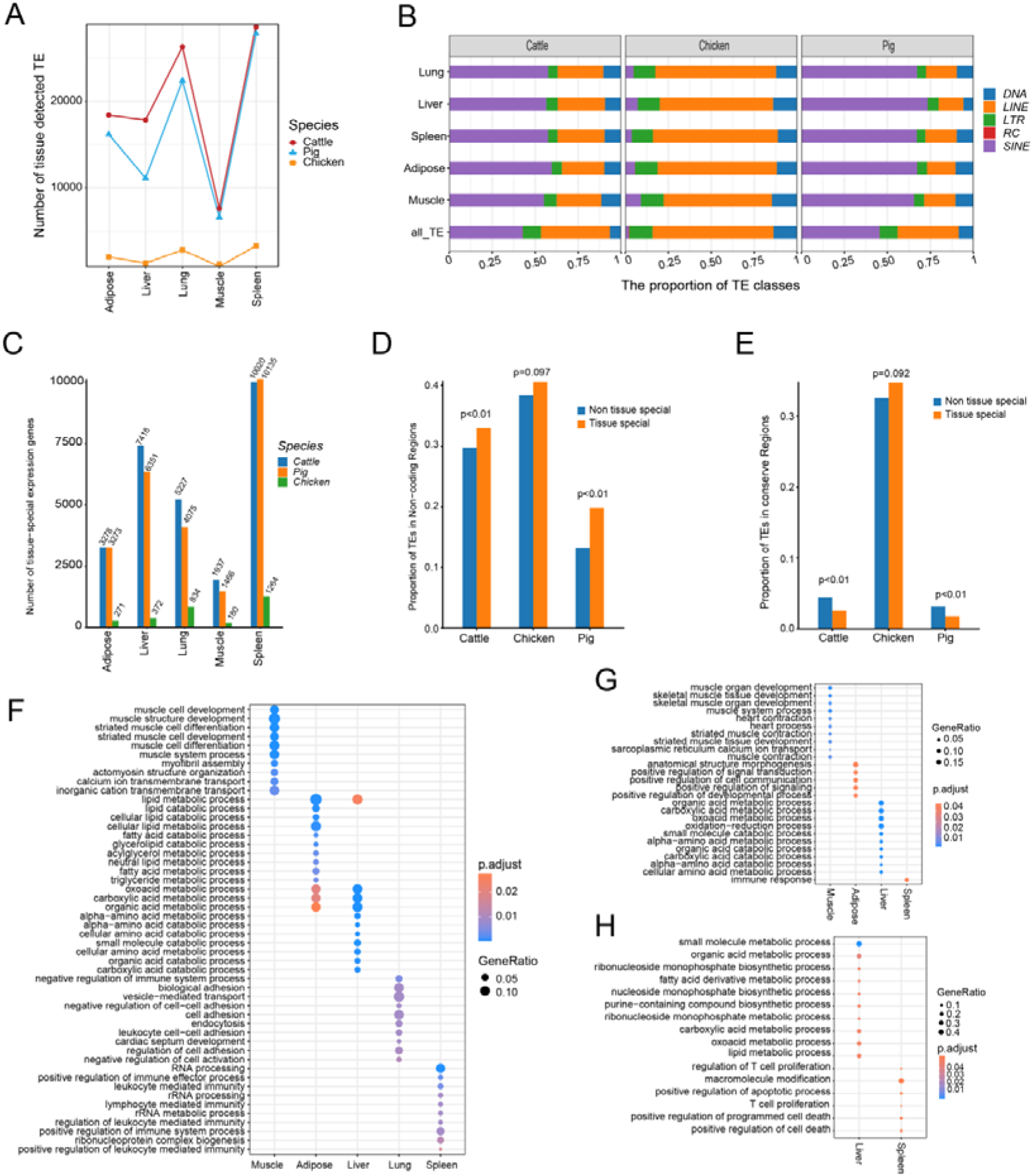
TEs specifically expressed in tissues are involved in tissue-related biological processes. (**A)** Dynamic changes in the number of detectable expressed TEs in different tissues of three species. (**B)** Comparison of the proportion of different classes of TEs among the detectable expressed TEs in each tissue (with the proportion of different classes of annotated TEs in the genome as background). (**C)** Number of tissue-specific expressed TEs in different tissues of three species. (**D)** The proportion of tissue-specific TEs and non-tissue-specific TEs in the non-coding region, p value is determined by Fisher’s exact test. E The proportion of tissue-specific TEs and non-tissue-specific TEs in the conserve element, p value is determined by Fisher’s exact test. (**F-H)** GO enrichment analysis of tissue-specific expressed TEs in pigs(**F**), cattle (**G**), and chickens (**H**) by extracting neighboring genes of tissue-specific expressed TEs. Only select the top 10 significantly enriched GO terms for tissue-specific expressed TEs for display.

As TE expression exhibited tissue specificity in different species by K-means clustering (Additional file 1: Fig. S5A-F), we further identified TEs detectable in a tissue-specific manner in the three species (Fig. 5C). Depending on their genomic insertion sites, TEs in the non-coding regions can potentially act as CREs to regulate gene expression. We performed enrichment analysis of tissue-specific TEs and non-tissue-specific TEs in the non-coding region of different species, and In pigs and cattle, tissue-specific TEs were significantly enriched in the non-coding region compared to non-tissue-specific TEs, while this difference was not significant in chickens (Fig. 5D). In addition, we found that non-tissue-specific TEs in pigs and cattle, but not in chickens, were more enriched in highly conserved DNA sequences compared to tissue-specific TEs (Fig. 5E). These findings suggest that, compared to those of chickens, TEs in the genomes of pigs and cattle made a more prominent contributions to tissue-specific CREs. In general, CREs specifically active in a certain tissue are known to play a crucial role in biological processes related to that tissues. In light of this knowledge, we investigated whether tissue-specific TEs also revealed such regulatory capacities. As TEs are known to have the potential to regulate the expression of their adjacent genes[49, 50], we performed gene ontology (GO) analysis on the neighboring genes of tissue-specific TEs (Additional file 4: Table S3A-C). We found that in pigs, neighboring genes of tissue-specific TEs in muscle, fat, liver, lung, and spleen were significantly enriched in tissue-relevant biological processes, respectively (Fig. 5F). Cattle had a similar enrichment pattern except for the lung (Fig. 5G). However, such an enrichment pattern was only observed in the liver and spleen in chickens (Fig. 5H).

Overall, our results suggest that TE expression exhibits tissue specificity in different species and that the tissue-specific expression of TEs implies regulatory potential in maintaining tissue-specific biological functions, especially in mammals.

### Using tissue-specific TEs to Dissect the GRNs underlying tissue-related biological processes

TE expression is regulated by chromatin modifications and TF binding [51]. Our study has revealed the biological significance of TEs that have tissue-specific expression or tissue-specific chromatin accessibility. Consequently, we proceeded to identify hub TEs in a tissue, which we defined as those exhibiting both tissue-specific expression and chromatin accessibility. To further study how tissue-specific TEs participate in tissue-relevant biological processes, we developed a computational framework to construct TE-GRNs. In contrast to conventional undirected GRNs, we leveraged these hub TEs to construct a directed GRN which encompasses transcriptional regulatory signaling from upstream TFs to TEs, then to downstream target genes. The specific strategy, illustrated in Fig. 6A, involves not only identifying TE-upstream TFs and TE-downstream target genes using a co-expression approach but also requires DNA sequence motif scanning to identify upstream TFs and examining TE-neighboring genes for potential downstream target genes.

**Figure 6.**
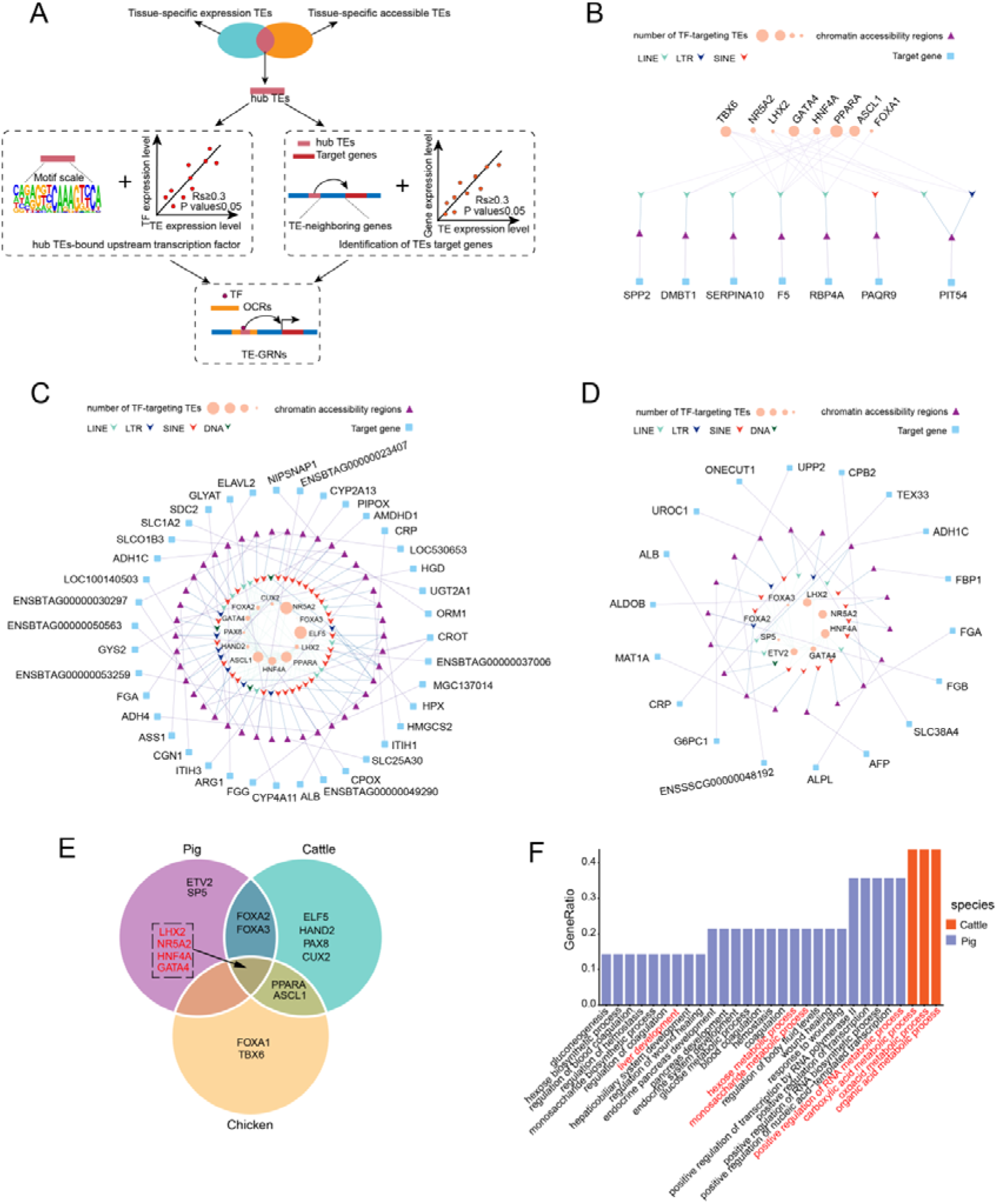
Tissue-specific TEs mediate gene regulatory networks involved in tissue-related biological processes. (**A)** The flowchart for constructing a gene regulatory network with tissue-specific TEs as the core. (**B-D)** TE-GRNs related to biological processes in liver tissues of chickens (**B**), pigs (**C**), and cattle (**D**) are shown. (**E)** The Venn diagram illustrates the specificity and sharing of upstream TFs for TEs in TE-GRNs related to the liver across the three species. The four TFs labeled in red and enclosed in the black dashed box are shared by the TE-GRNs of the three species. (**F)** GO enrichment analysis on the downstream target genes of TEs in pig and cattle TE-GRNs. The GO terms highlighted in red are closely associated with liver development or metabolism.

Since the liver is an important metabolic organ and substantially affects many economic traits of domesticated animals [52–54], we used it as an example to demonstrate the construction of TE-GRNs underlying liver-relevant biological processes in pigs, cattle, and chickens (Fig. 6B-D, Additional file 5: Table S4A-C). Analyzing TE-GRNs within the liver across the three species yielded insights into the shared and species-specific TFs upstream of hub TEs. For example, we identified four TE-upstream TFs (LHX2, NR5A2, HNF4A, and GATA4) that were common to all three species. Additionally, we discovered two TFs (ETV2 and SP5) specific to pigs, and four TFs (ELF5, HAND2, PAX8, and CUX2) specific to cattle. Chickens exhibited two species-specific TFs (FOXA1 and TBX6) (Fig. 6E). Notably, among the identified upstream TFs associated with these hub TEs in the three species, we observed a significant number of TFs that exhibited a remarkable enrichment of motifs within TEs with liver-specific chromatin accessibility, including HNF4A, the FOXA gene family, and PPARA (Fig. 3E, Additional file 1: Fig. S2C-D). HNF4A[55, 56], GATA4[57, 58], FOXA gene family [59, 60], and PPARA[61]have been reported to play important regulatory roles in liver-relevant biological processes. These findings support the authenticity and reliability of our approach in identifying key TFs acting upstream of TEs. To further confirm that the liver TE-GRNs are involved in important biological processes of the liver, we conducted GO analysis of the genes in these networks of the three species, respectively. Genes in TE-GRNs of pigs and cattle are significantly enriched in metabolic pathways, and genes in pig TE-GRNs are also enriched in the biological pathways of liver development (Fig. 6F). However, genes in chicken TE-GRN did not enrich in liver related biological processes. This could potentially be attributed to the small number of genes in chicken TE-GRN and a correspondingly lower statistical power for enrichment analysis. Nevertheless, among the TE-downstream target genes in chicken TE-GRN, several genes are closely related to metabolism, including PAQR9, which can influence hepatic ketogenesis and fatty acid oxidation by regulating the protein stability of PPARA[62]. Additionally, PIT54 is highly expressed in the chicken liver and may be closely associated with liver physiology, function, and development[63].

In summary, we presented a computational framework to decipher TE-GRNs underlying liver-related biological processes. Our strategy not only facilitates future understanding of the genetic basis of liver-related economic traits, such as feed conversion rate and lipid metabolism, but also holds great promises for dissecting the molecular underpinnings of other important economic traits by adopting a TE-centric perspective.

## Discussion

TEs are a vital component of the genome that can influence diseases or phenotypes by modulating gene regulatory networks. With the growing recognition of the functional significance of TEs, an expanding body of research has embraced the integration of multi-omics data. This integration has enabled investigations into the regulatory role of TEs in a diverse range of biological processes, not only in humans but also in model animals. Nevertheless, the livestock genetics community has been lagging behind in generating such data, leading to a shortage of research that comprehensively explore the regulatory landscape of TEs in domesticated animals.

In our research, we constructed comprehensive maps of TEs in pigs, cattle, and chickens, and compared their evolutionary dynamics as well as contributions to different types of CREs. We systematically analyzed the expression of TEs across tissues, revealing the importance of tissue-specific TEs in maintaining tissue-specific biological processes. Given the important role of TEs in rewiring transcriptional regulatory networks, we presented a novel computational framework to constructTE-GRNs based on tissue-specific TEs. As a proof of concept, we built TE-GRNs in the liver tissues of pigs, cattle, and chickens, identifying TE-upstream TFs and TE-downstream target genes that gain multiple lines of literature support. To our best knowledge, our study is the first to explore the TE regulatory landscape of multiple tissues in the three major domesticated animals, and presented a computational framework that facilitates studying how TEs affect molecular pathways underlying complex traits and diseases.

The genome annotation of TEs is essential for the study of any TE-related biological questions. Following the TE annotation efforts in several previous studies[34–37], we used ReatMacker software to annotate TEs in the reference genomes of pigs, cattle, and chickens, respectively. Our analysis showed that TEs in pigs and cattle accounted for 43.4% and 47.5% of their respective genomes, while in chickens, TEs made up only 9.6% of the genome. Given that propagation of TE copies during evolution have expanded the genome size, previous studies have suggested a positive correlation between the level of TE proportion and the size of a genome [43, 44]. Consistently, this correlation holds true for TE proportions across the genomes of pigs, cattle and chickens. Furthermore, we identified TE families that are species-specific, such as SINE/tRNA in pigs, LINE/CR1 in chickens, and SINE/Core-RTE, SINE/tRNA-Core-RTE, and LINE/RTE_BovB in cattle. Our analysis also confirms several previous studies, which showed that Chicken Repeat 1 (CR1) was the most abundant repeat family in chickens[64], SINE/tRNA was the most abundant repeat family in pigs [65], and non-LTR LINE RTE (BovB) and BovB-derived SINEs were cattle-specific repeats [66].

In this study, we analyzed the age distribution of all annotated TEs in three species based on the single nucleotide mutation rate in each species’ genome. By comparing the burst patterns of different TE classes and families during evolution, we identified TE types that still exhibit transposition activity in the modern genome of specific species. For instance, SINE/tRNA and LINE/L1 have undergone high levels of amplification during pig evolution and remain active in the modern pig genome. Notably, a recent burst in SINE/tRNA was observed, consistent with previous findings that SINE/tRNA is currently the most active transposon type in the modern pig genome [67–69]. In this work, we also analyzed the relationship between the genomic distribution density of different classes of TEs in the three species and found that the correlation levels of genomic distribution density between different TEs have species-specific and shared pattern. For example, LTR and LINE densities are positively correlated in all three species. SINE density is negatively correlated with LINE densities only in pigs and cattle, which is consistent with previous reports on mammals [36, 70]. Additionally, SINE density shows a stronger negative correlation with LTR densities, while LINE density is positively correlated with LTR and DNA densities in chickens. These findings suggest that the modes of cooperation among different classes of TEs in maintaining genome stability may vary across species.

In the current study, we also systematically investigated the impact of TEs on epigenetic marks associated with CREs. Our findings showed that TEs have a greater contribution to the OCRs in pigs and cattle than in chickens (Fig. 3B). This difference might imply the differential contributions of TEs to CREs between mammals and avians. Previous studies have demonstrated that TEs can act as CREs to regulate gene expression by carrying specific TF binding motifs [71–76]. Similarly to He et al. [77], we found that tissue-specific TEs enriched in pigs, cattle, and chickens contain numerous TF binding motifs related to tissue growth and development, suggesting that tissue-specific TEs are crucial for maintaining tissue-related biological processes. A genome contains diverse types of CREs that regulate gene expression through different molecular mechanisms. To further explore the similarities and differences in the effects of TEs on the various types of CREs in the three species, we identified promoters, enhancers, and CSEs in pigs, cattle, and chickens, and built a multitask deep learning model that showed good performance in classifying the three CRE types by DNA sequences. We systematically analyzed the differences in the extent to which TEs contributed to CREs in the three species, which enhanced our understanding of the differential potential of TEs to influence CREs in different species.

In addition to their primary role in transposition, TEs exhibit a multifaceted impact on biological systems, influencing various aspects such as gene expression, chromatin accessibility, activation of cellular signaling pathways (including the interferon response), RNA interference responses, and even contributing to aging processes or antiviral activities [78]. Genes expressed in a tissue-specific manner are intricately linked to the tissue-relevant biological processes. While there are several sources available for tissue-specific gene expression, including HPA [79], TiGER [80], GTEx [81]and TissGDB [82], research on tissue-specific TE expression is still limited. In this study, we not only constructed TE expression maps for five tissues in the pigs, cattle and chickens for the first time, but also identified TEs with tissue-specific expression patterns. We showed that these tissue-specific TEs are intimately associated with tissue-related biological processes. Therefore, our study provides a valuable reference for future investigations on how TEs mediate epigenetic regulation in the evolution and domestication of pigs, cattle, and chickens.

To further dissect how TEs regulate various biological processes, we developed a computational framework to construct TE-GRNs. In the liver tissue, the TE-GRNs consisted of TE-downstream target genes that are enriched in liver-related biological processes, as well as TE-upstream TFs that include liver-related pioneering TFs, which supports the reliability of our network construction approach. While our TE-GRN is a promising advancement in understanding TE biology, there is still considerable room for improvement in its accuracy and comprehensiveness. With the increasing availability of different types of multi-omics data in various biological contexts, identifying TE-downstream target genes can be improved by chromatin looping information from HiC data, and identifying TE-upstream TFs can be facilitated by high-throughput protein-DNA interaction data, such as ChIP-Seq and CUT & Tag. With the increasing generation of scRNA-seq and scATAC-seq data across diverse biological contexts and the ongoing development of quantitative tools for assessing single-cell TE expression [77, 83], the TE-GRN construction approach could be adopted to studying how TEs contributed to transcriptional regulatory networks at cell type-specific level. This will enable more powerful analyses of the TE-mediated phenotype variations that exclusively manifest within certain cell types. Overall, By revealing the molecular pathways involved in tissue-related biological processes, the development of TE-GRNs represents a valuable resource for future investigations into the molecular basis of phenotypic variation, from a TE-centric perspective.

## Conclusions

In this work, we comprehensively analyze TEs in three major domesticated animals: pigs, cattle, and chickens, exploring their potential roles in genomic evolution, epigenetic mechanisms, and gene expression regulation. In addition, we present a computational framework for constructing TE-GRNs, providing a better understanding of the impact of TEs on complex traits and diseases. Our work significantly contributes to our understanding of the importance of TEs and their potential roles in shaping the evolutionary trajectory and deciphering complex traits of domesticated animals.

## Methods

### Data Sources

In this study, all data were sourced from public databases. To annotate TEs, we utilized reference genome sequences (bosTau9.fa, galGal6.fa, susScr11.fa) for cattle, pigs, and chickens obtained from the UCSC database. For assessing the potential functionality of TEs, we obtained ATAC-seq, Chip-seq, and RNA-seq data for liver, spleen, lungs, muscles, and adipose tissues of pigs, cattle, and chickens from a publication by Zhou et al. (GSE158430) [39]. To ensure a conservative analysis, we obtained highly conserved element regions specific to each species (gerp_constrained_elements.bos_taurus.bb, gerp_constrained_elements.gallus_gallus.bb, gerp_constrained_elements.sus_scrofa.bb) from the Ensemble database.

### TE annotation

In this study, TE annotation was performed using RepeatMasker [84] software (version 4.1.2) with the rmblastn engine, along with the Dfam_Consensus-20181026 and Repbase-20181026 libraries. The following parameters were set: -species, -a, -s, -gff, -cutoff 225. For calculating TE divergence and assessing the coverage of different TE families in the genome, we utilized calcDivergenceFromAlign.pl and createRepeatLandscape.pl, respectively. To estimate the relative age of TEs in the genome, we employed the TE divergence values obtained from RepeatMasker software, applying the formula T = K / 2r [85]. For TE age calculations, we used a consistent single nucleotide mutation rate of r = 2.2 × 10^−9^ substitutions/site/year across mammalian genomes [86] and a mutation rate of 1.91 × 10^−9^ substitutions per site per year in avian genomes [87]. Consequently, we set r as 2.2 × 10^−9^ for pig and cattle TE age calculations, and 1.91 × 10^−9^ for chicken TE age calculations in this study.

### TE genomic landscape analysis

In this study, we used the “createRepeatLandscape.pl” command to extract the genome coverage of different TE types. The proportion of each class or family was calculated by dividing its count by the total number of annotated TEs. To annotate the genomic distribution of TEs, we employed the R package ChIPseeker[88] and utilized genome annotation files from the Ensemble database (Sus_scrofa.Sscrofa11.1.100.chr.gff3, Bos_taurus.ARS-UCD1.2.104.gff3, Gallus_gallus.GRCg6a.104.chr.gff3). For investigating the distribution of TE age across the three species, we determined the count of each class or family at different time points based on the insertion age of each TE. All statistical results were visualized using the R package ggplot2 [89]. In exploring the relationship between the genomic distribution of different TE classes, we followed a similar approach as previous studies [36].

### Analyzing the impact of TEs on OCRs

To assess the impact of TEs of different ages and types on OCRs in the three species, we performed an integrated analysis with ATAC-seq data to examine the relationship between TEs and chromatin accessibility. Initially, we obtained raw ATAC-seq data and conducted quality control using Trim Galore. The reads were aligned to the reference genome using Bowtie2 [90]. We then removed PCR duplicates using Sambamba [91] and converted BAM files to BigWig files with deepTools[92]’s bamCoverage script (--extendReads --normalizeUsing RPKM). Next, we quantified the chromatin accessibility signals of different TE classes in different tissues based on the BigWig files using the computeMatrix command in deepTools. Finally, we visualized the chromatin accessibility signals of TEs in different tissues of the three species using the plotProfile command.

For OCRs, we obtained peak files from five different tissues (with two biological replicates per tissue) for pigs, cattle, and chickens. We identified consistently present OCRs in each tissue by utilizing the intersect and merge commands in BEDTools [93]. To investigate the direct impact of TEs on OCRs, we used the intersect command in BEDTools to calculate the proportion of OCRs affected by TE insertions, referred to as TE-driven OCRs. We categorized TEs into age groups: youngest TEs (0-25 million years (Mya)), younger TEs (25-50 Mya), older TEs (50-75 Mya), and oldest TEs (75-100 Mya). Subsequently, we calculated the proportions of different classes and ages of TE-driven OCRs in each tissue of the three species, with proportions of all genome-annotated TE classes and ages as background. All statistical results were visualized using the ggplot2 package in R.

To identify TEs with chromatin accessibility in specific tissues, we utilized the intersectBed function of BEDTools with parameters of -f 0.5 and -F 0.2 to determine overlaps between TEs and OCR peaks in each tissue [94]. We considered a TE to be associated with chromatin accessibility if it overlapped with OCR peaks by at least 50%, and OCR peaks overlapped with TEs by at least 20%. Shared and tissue-specific TE accessibility was visualized using the UpSetR package [95] in R, based on the TEs identified with chromatin accessibility in each tissue. Tissue-specific accessible TEs were identified using the setdiff function in R. To identify enriched TF motifs in tissue-specific accessible TEs, we performed motif enrichment analysis using HOMER software [96] on tissue-specific TEs in each tissue. We selected the top 20 significantly enriched TFs that were expressed in the corresponding tissue for each species. Bubble plots illustrating the results were generated using the ggplot2 package in R.

### Identification and Deep Learning Classification of different DNA CREs

To investigate the effects of different TE classes and families on DNA CREs in various species and tissues, we downloaded CTCF, H3K4me1, H3K27ac, and H3K4me3 peak region files for five tissues of pigs, cattle, and chickens, each with two biological replicates. To obtain consistent peak regions across tissues, we merged the biological replicates using the BEDTools intersect and merge commands. Based on the consistent histone modification peaks across tissues, we identified enhancer, promoter, and putative silencer regions for each tissue in different species. For enhancer regions, we defined H3K27ac signal regions located outside of known coding gene TSS ± 1kb and without overlap with H3K4me3 peaks. Promoter regions were defined as H3K27ac signal regions overlapping with known coding gene TSS ± 1kb or H3K4me3 peaks. Putative silencer regions were identified following a previously described method [97]. In this method, chromatin open regions overlapping with CTCF, H3K4me1, H3K27ac, H3K4me3, and TSS were excluded, and the remaining chromatin open regions were defined as candidate silencer regulatory elements.

To assess the reliability of the identified CSEs and examine sequence feature differences among the three types of regulatory elements, we constructed a multitask deep learning classification model based on Almeida’s Deepstarr regression model [98]. Within ±1kb of the center point of each regulatory element, we extracted DNA sequence information and converted it into one-hot encoding format as input data for the deep learning model. The sequence labels were also converted into one-hot encoding format based on different regulatory element types. The input data were shuffled and split, with 15% reserved for the test set, and the remaining data used for model training (80%) and validation (20%) (Additional file 6: Table S5A-C). To evaluate the performance of each regulatory element type classification accuracy, we used the R package plotROC [99] to calculate and visualize the AUC results based on the predicted and true values of the test set.

### Contribution of TEs to different DNA CREs

To investigate the extent to which different types of CREs (Promoters, Enhancers, CSEs) are influenced by TEs, we utilized the “intersect” command in BEDTools to extract TE sequences from specific types of DNA CREs. We then quantified the proportion of TE bases relative to the total bases within each regulatory element using R language and used the genomic coverage of all annotated TEs in the species as a reference. By this method, we systematically revealed the extent to which TEs influence the composition of different types of CREs in different tissues of the three species. To investigate the contribution of different classes of TEs to the composition of CREs, we quantified the proportion of TE bases for each TE classes within the CREs, In this part of the analysis, the proportion of TE base for each type of TE in all annotated TEs was used as the background.

In addition, to investigate the contribution of TE insertions to different types of CREs, we utilized the ‘intersect’ command in BEDTools to calculate the proportion of annotated TEs overlapping with specific regulatory element across tissues in three species. To further focus on the ability of different classes of TEs to integrate into different CREs, we explored the proportion of different classes of TEs based on those TEs overlapping with specific regulatory element. The above statistical results were visualized using the ggplot2 R package in R language.

### Enrichment analysis of TE families in different DNA CREs

To investigate the enrichment of TE families in different CREs across different tissues of the three species, we employed a permutation test, randomly sampling size-matched regulatory element regions in the genome 1000 times to calculate the fold enrichment and significance of different TE families in each type of regulatory element. The significance P-values were corrected using the Bonferroni method. Prior to conducting the enrichment analysis, we filtered out low-abundance TE families since they may have low statistical power due to their low copy number in certain species. In this study, TE families with more than 3000 copies in pigs and cattle, and more than 1000 copies in chickens were retained for analysis. The results of the enrichment analysis were integrated using R language and visualized using the pheatmap package in R.

### Quantification of TE loci expression and identification of tissue-specific TE expression

To detect the expression activity of TEs in each tissue of three species, we downloaded transcriptome data of liver, spleen, lung, muscle, and adipose tissue from three species, with two biological replicates for each tissue. First, we conducted quality control on the transcriptome data of each sample using Trim Galore software (-q 20 --phred33 --stringency 4 --length 20 -e 0.1). Then, based on the annotation information of TEs in the species genomes and RNA-seq clean data, we used SQuIRE software to detect the transcriptional activity of TEs in each sample. To remove the interference of low-expression TEs on downstream analysis, we defined TEs with an average FPKM≥1 in the two biological replicates of a tissue as detectable TEs in that tissue. By integrating the expression levels of detectable TEs in the five tissues and using the R package ComplexHeatmap [100] for plotting K-means heatmaps. To identify tissue-specifically expressed TEs, based on the matrix formed by the expression values of detectable TEs in each sample, after data normalization, we calculated the tissue-specificity index (TSI) of each TE using the following formula: 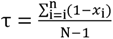 [101]

Tissue-specifically expressed TEs were identified as those with a delta value greater than 0.9. TEs with a normalized expression value of 1 within a tissue were considered specifically expressed in that tissue.

### Enrichment analysis of tissue-specific expressed TEs

To explore the enriched TE classes in the expressed TEs detected in each tissue, we compared the proportion of different TE classes in the detectable TEs to their proportion in all annotated TEs in the species, serving as the background. This allowed us to identify any significant changes in the proportion of TE classes, and to observe any enrichment or depletion of certain TE classes in the expressed TEs of each tissue compared to the background.

To investigate the enrichment of tissue-specifically expressed TEs in genomic coding regions and conserved elements, we obtained conserved element information for the genomes of three species from Ensembl and defined genomic coding regions based on genome annotation files, with gene body regions of protein-coding genes considered as coding regions. Non-tissue-specifically expressed TEs were used as controls, and we tallied the numbers of tissue-specific and non-specific TEs in the coding regions and conserved elements. Contingency tables were constructed, and we conducted Fisher’s statistical testing using the fisher.test function in R. The results were visualized using the R package ggplot2.

To examine the biological processes associated with tissue-specific TE expression, we annotated tissue-specific TEs’ genomic distribution using ChIPseeker. We extracted neighboring genes of TEs as potential target genes and performed GO function enrichment analysis with the R package clusterProfiler[102]. For result visualization, we selected the top 10 significantly enriched GO terms in each tissue and used the R package ggplot2.

### Construction of TE-GRNs

Since tissue-specific TEs are involved in tissue-related biological processes, we focused on constructing directional gene regulatory networks with TEs as the central. Our first step was to identify these hub TEs by examining those that overlap with tissue-specific OCRs and exhibit tissue-specific expression patterns. We then determined the upstream TFs and downstream target genes regulated by these hub TEs. For identifying the downstream target genes of hub TEs, we adopted a strategy inspired by previous studies [83, 103, 104] that suggest the transcriptional activity of TEs may potentially influence the expression of neighboring genes and the co-expression patterns between TEs and their neighboring genes. To perform co-expression analysis between TEs and genes, we quantified the expression of genes in each transcriptome sample as follows: We performed genome alignment of the transcriptome data using Hisat2 software [105] (--rna-strandness RF), followed by quantification of gene reads using the featureCounts software [106] (-s 2). Finally, the expression level of each gene was quantified by calculating TPM values. Next, to identify the downstream target genes of hub TEs, we performed genomic annotation of hub TEs using the ChIPseeker R package, obtaining their neighboring genes. Subsequently, we assessed the linear regression relationship between TE and neighboring gene expression levels in different tissues using the lm function in R. Statistically significant TE-gene pairs (model P value < 0.05) were filtered, and the Spearman correlation between TE-gene pairs was calculated using the R base function cor.test to determine positive regulatory relationships (Rs ≥ 0.3). To screen for the upstream TFs of hub TEs, we employed a multi-criteria screening strategy. Firstly, we used the BEDtools getfasta command to accurately extract TE DNA sequences from the genome based on their strand information, confirming the presence of TF binding sites in TE sequences. Then, we downloaded the Custom Motif Matrices files of TFs from http://homer.ucsd.edu/homer/custom.motifs and utilized the scanMotifGenomeWide.pl command in HOMER software to scan for motifs within TE sequences and identify TF-TE pairs. Subsequently, we used the R base function cor.test to calculate the correlation between TF-TE pairs (method = “spearman”, adjust = “fdr”). TF-TE pairs with Rs ≥ 0.3 and adjust.p < 0.05 were considered to have a regulatory relationship. Additionally, by identifying the co-localization of hub TEs and CREs, we were able to infer the CREs mediated by hub TEs. Overall, we determined the regulatory relationships between TF binding to TEs, resulting in chromatin opening and the regulation of target genes. Finally, we visualized these regulatory relationships in liver tissue using the Cytoscape [107] software.

## Additional files

**Additional file 1**

Format: .docx

Title: Additional file 1 Figure S1-5.

Description: Supplementary Figure 1-5.

**Additional file 2**

Format: .xlsx

Title: Additional file 2 Table S1.

Description: Supplementary Table 1.

**Additional file 3**

Format: .xlsx

Title: Additional file 3 Table S2.

Description: Supplementary Table 2.

**Additional file 4**

Format: .xlsx

Title: Additional file 4 Table S3.

Description: Supplementary Table 3.

**Additional file 5**

Format: .xlsx

Title: Additional file 5 Table S4.

Description: Supplementary Table 4.

**Additional file 6**

Format: .xlsx

Title: Additional file 6 Table S5.

Description: Supplementary Table 5.

