## Additional file 1 for "Multi-omics analysis reveals critical cis-regulatory roles of transposable elements in livestock genomes"

**Figure S1. Genomic distribution characteristics of different TE types.** (**A-C**) The genome distribution of diffirent TE families of pig (**A**), cattle (**B**), and chicken (**C**). (**D and E**) Correlation between the genomic distribution density of different TE classes in the cattle (**D**) and chicken (**E**) genome.

**Figure S2. Tissue-specific accessibility analysis of TEs.** (**A-B**) Upset plot showing tissue-specific accessibility of TEs in cattle (**A**) and chicken (**B**). (**C-D**) Enriched transcription factor (TF) motifs in five tissue-specific accessible TEs of cattle (**C**) and chicken (**D**).

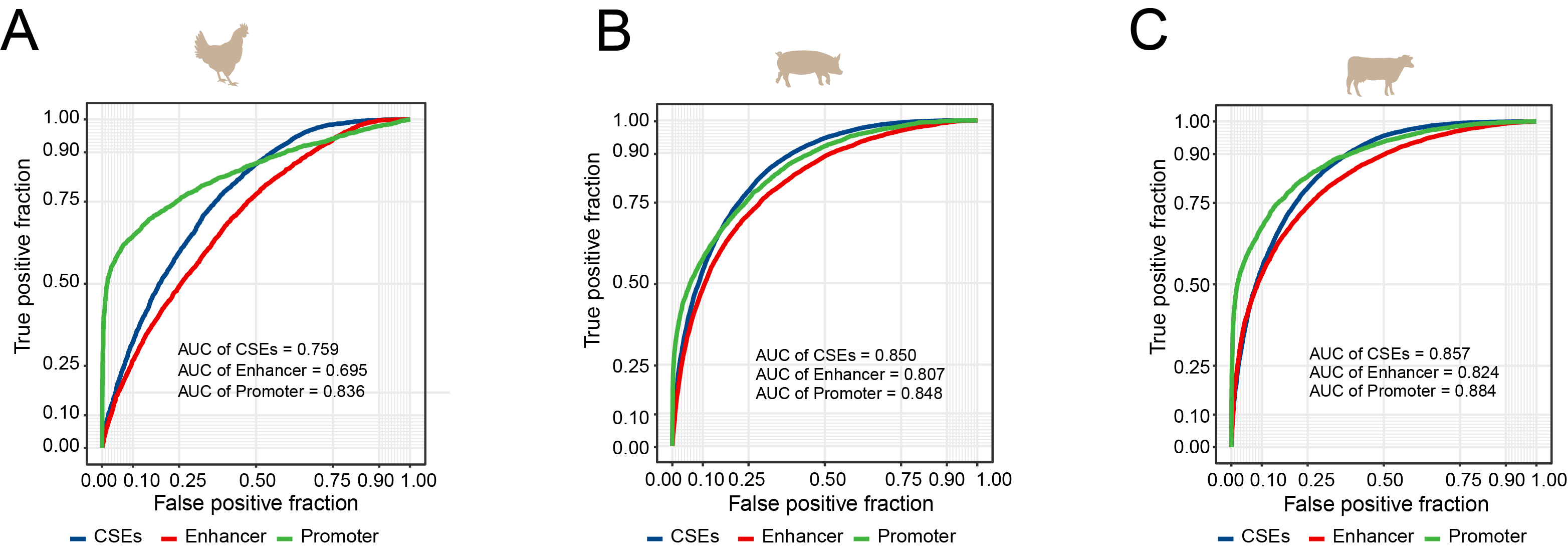

**Figure S3. Accurate Classification of CREs using Multi-Task Deep Learning Model.** (**A-C**) The AUC plot shows the accuracy of the deep learning model in classifying three types of elements in chickens (**A**), pigs (**B**), cattle (**C**).

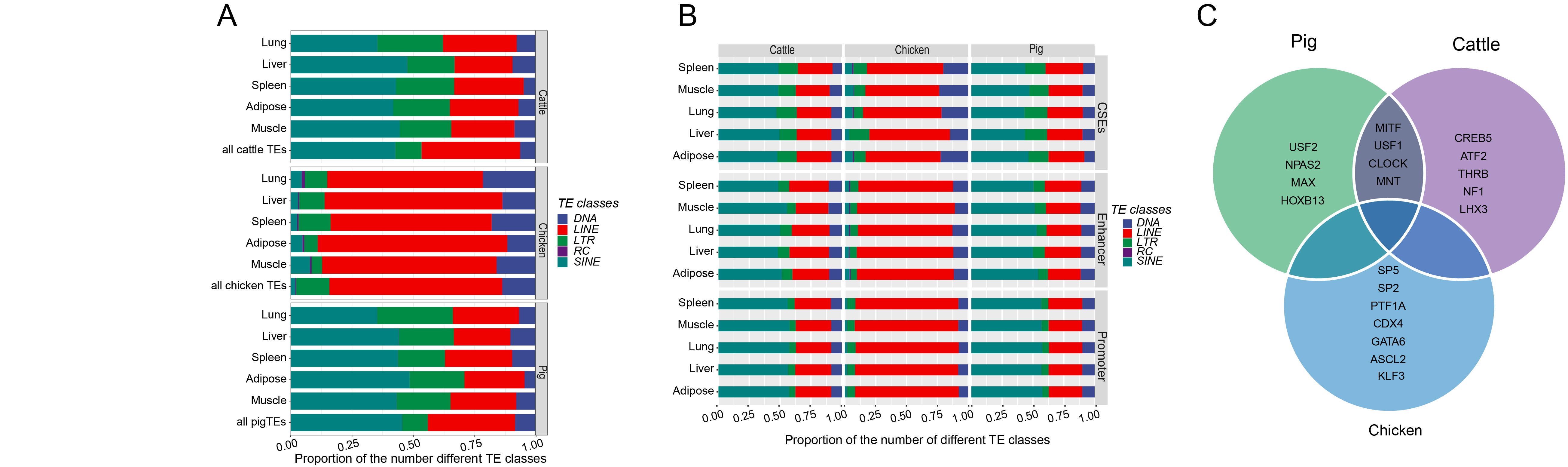

**Figure S4. Contribution of TEs to CRE activity.** (**A**) The proportion of different TE classes within CREs, with the background proportion of different TE classes in all annotated TEs of the species. (**B**) The proportion of different TE classes in specific-regulatory element state TEs across different species and tissues. (**C**) Venn diagrams displaying TFs corresponding to DNA motifs significantly enriched in LTRs within CSEs for each species. Only the top 20 enriched TFs, and belonging to genes in that species, are shown.

**Figure S5. Cluster analysis of heatmaps of detectable expressed TEs in Tissues.** (**A-C**) WSS plot of detectable TEs in pigs (**A**), cattle (**B**), and chickens (**C**) to determine the number of clusters. (**D-F**) Heatmap showing dynamic expression of detectable TEs in pigs (**D**), cattle (**E**), and chickens (**F**) across tissues.
